# Tuberculosis outbreak investigation using phylodynamic analysis

**DOI:** 10.1101/229500

**Authors:** Denise Küehnert, Mireia Coscolla, David Stucki, John Metcalfe, Lukas Fenner, Sebastien Gagneux, Tanja Stadler

## Abstract

The fast evolution of pathogenic viruses has allowed for the development of phylodynamic approaches that extract information about the epidemiological characteristics of viral genomes. Thanks to advances in whole genome sequencing, they can be applied to slowly evolving bacterial pathogens like *Mycobacterium tuberculosis*.

In this study, we investigate the epidemiological dynamics underlying two *M. tuberculosis* outbreaks using phylodynamic methods. The first outbreak occurred in the Swiss city of Bern (1993-2012) and was caused by a drug-susceptible strain belonging to the phylogenetic *M. tuberculosis* Lineage 4. The second outbreak was caused by a multidrug-resistant (MDR) strain of Lineage 2, imported from the Wat Tham Krabok (WTK) refugee camp in Thailand into California.

There is little temporal signal in the Bern data set and moderate temporal signal in the WTK data set. We estimate an evolutionary rate of 0.0039 per single nucleotide polymorphism (SNP) per year for Bern and 0.0024 per SNP per year for WTK. Nevertheless, due to its high sampling proportion (90%) the Bern outbreak allows robust estimation of epidemiological parameters despite the poor temporal signal. Conversely, there’s much uncertainty in the epidemiological estimates concerning the WTK outbreak, which has a small sampling proportion (9%). Our results suggest that both outbreaks peaked around 1990, although the Bernese outbreak was only detected in 1993, and the WTK outbreak around 2004. Furthermore, individuals were infected for a significantly longer period (around 9 years) in the WTK outbreak than in the Bern outbreak (4-5 years).

Our work highlights both the limitations and opportunities of phylodynamic analysis of outbreaks involving slowly evolving pathogens: (i) estimation of the evolutionary rate is difficult on outbreak time scales and (ii) a high sampling proportion allows quantification of the age of the outbreak based on the sampling times, and thus allows for robust estimation of epidemiological parameters.

## Introduction

Whole genome sequencing (WGS) of clinical *M. tuberculosis* isolates is performed retrospectively and allows to confirm/refute suspected epidemiological links to identify individuals contributing to transmission, and explore drug-resistance and/or compensatory mechanisms which emerged during anti-tuberculosis treatment [1–6]. Although WGS analysis has the potential to reveal more complex epidemiological dynamics such as how long the outbreak was not controlled, the time patients are infectious for, the proportion of sampled cases, and the transmission potential of different strains, these epidemiological parameters are rarely estimated for slowly evolving bacterial pathogens such as *M. tuberculosis* [2, 5, 7]. In the context of tuberculosis disease, answering those questions may help to evaluate and improve treatment strategies and control programs.

Phylodynamic analysis of real time WGS data can shed light on temporal dynamics of disease outbreaks, for example to determine if there is ongoing transmission [5]. Here, we employ phylodynamic methods to shed further light on two *M. tuberculosis* outbreaks. The first outbreak was detected around 1991 in the city of Bern, Switzerland, where twenty-two related cases, mainly homeless individuals and substance abusers, were identified initially [8]. Using a novel combination of strain-specific SNP screening assay and targeted WGS, a tuberculosis cluster spanning 21 years and involving 68 patients was identified [3, 7]. The genomic analysis revealed that this outbreak was caused by a Lineage 4 strain (Euro-American) of *M. tuberculosis*, and all but one showed no evidence of antibiotic resistant conferring mutations. The analysis revealed three sub-clusters within the outbreak, one of them associated to HIV coinfection.

The second data set consists of 30 MDR strains imported to California during resettlement of refugees from the refugee camp at Wat Tham Krabok (WTK) [4]. Whole genome analysis confirmed that the strains causing the outbreak were multidrug-resistant and belonged to the Lineage 2 (East-Asian, Beijing genotype) of *M. tuberculosis*. Genomic data supported a single case whose isolate occupied the central node of the transmission network indicating the presence of a super-spreader. Epidemiological data integrated with the transmission chain also demonstrated multiple independent importation events from Thailand with reactivation and transmission within California over a 22-year period.

In this study, we aim to understand the dynamics of tuberculosis outbreaks by inferring phylogenetic trees together with epidemiological parameters, in particular, transmission and recovery rates, from genome sequence data using phylodynamic methods.

## Methods

### Reconstruction of transmission dynamics

First, we explored the temporal signal in the sequence alignments using TempEst [9].

The main analysis of both data sets was done within the Bayesian MCMC framework BEAST2 [10]. We assume that the phylogeny spanned by the genomic samples is a suitable approximation of the transmission tree, such that we can estimate epidemiological parameters simultaneously with the phylogenetic tree. We employ two phylodynamic methods, the birth-death skyline plot (BDSKY) [11] and the multi-type birth-death model (MTBD) [12]. Both assume that an infection event can be considered as the "birth" of a newly infected individual, while a recovery event (successful treatment) is a "death". While the BDSKY model assumes that an infected individual is immediately infectious upon infection, the MTBD model allows us to incorporate the fact that *M. tuberculosis* infections usually start with a latent period in which the infected individual is not yet infectious.

In both analyses we employ a general time reversible substitution model with gamma distributed rate heterogeneity and a proportion of invariant sites (GTR + I + Γ). A relaxed lognormal clock is used to model the variation of evolutionary (substitution) rates across branches, such that we estimate a mean clock rate θ (per SNP per year) and standard deviation σ for the lognormal branch rate distribution. All parameters are estimated jointly. The prior distributions used are summarized in **Table 1**.

**Table 1.**
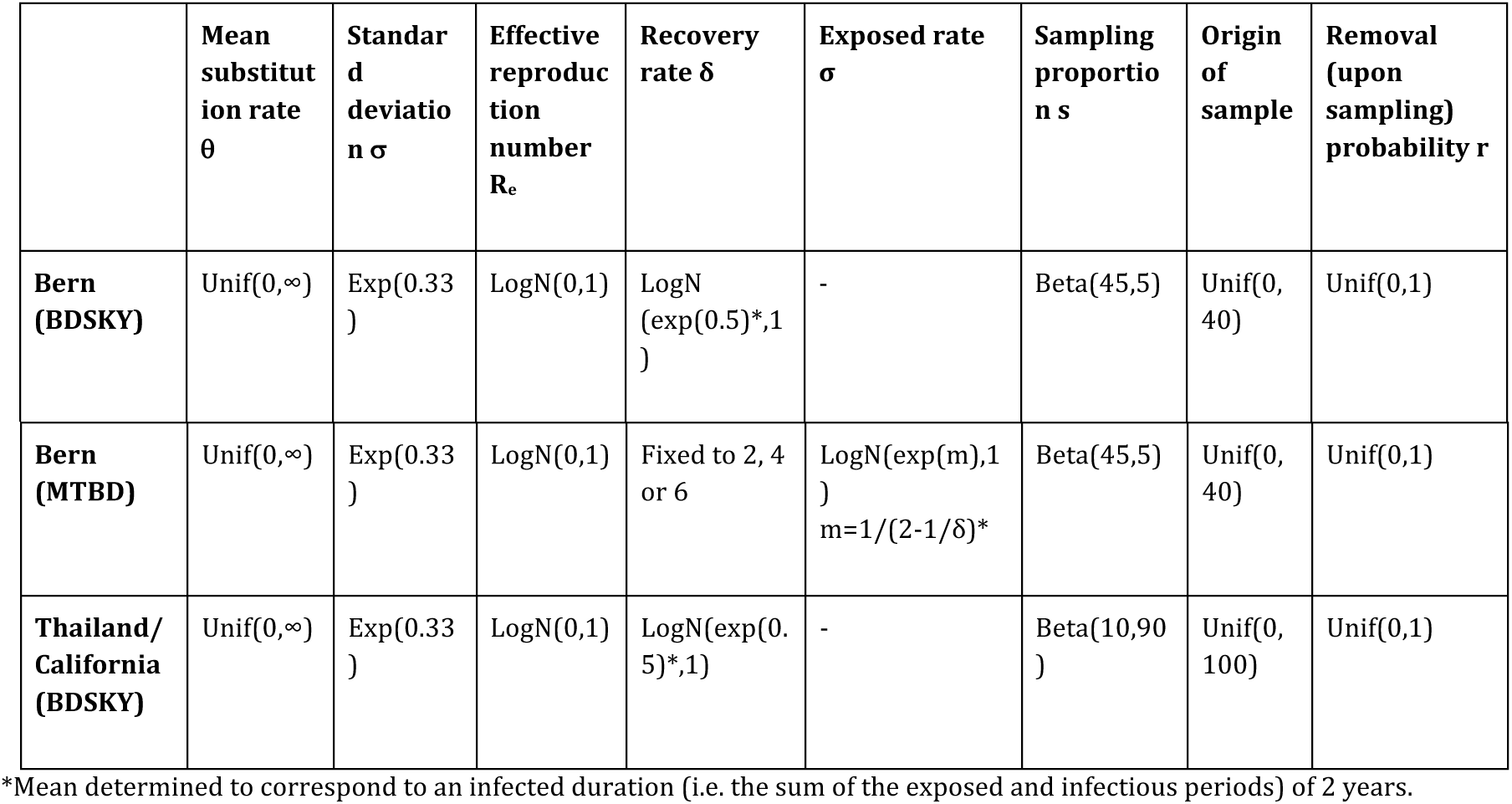

#### Phylodynamic analysis with the birth-death skyline model

The birth-death skyline model [11] describes a prior distribution for a transmission tree and is based on a stochastic birth-death process, with birth (λ), death (μ) and sampling (ψ) rates. Individuals become non-infectious upon sampling with probability r ∈ [0,1] [13]. Typically, the probability r is close to 1 if sampling is accompanied by successful treatment. To investigate the change of epidemiological dynamics, the period covered by the phylogeny is divided into intervals, and parameters are constant within an interval but may change between intervals. We can estimate the effective reproduction number R_e_, through the alternative parametrization of the model using the effective reproduction number R_e_ = λ /(μ + rψ), the rate at which individuals become non-infectious δ = μ + rψ and the sampling proportion s = ψ / (μ + ψ). We employ m = 5 intervals to estimate potential changes in R_e_, and assume that δ is constant through time. The sampling proportion s is set to zero before the first sample, and assumed to be a positive constant thereafter.

#### Phylodynamic analysis with the multi-type birth-death model – incorporating the latent period

The MTBD model allows us to incorporate and investigate the exposed phase. In the following we use the terms ‘latent’ and ‘exposed’ interchangeably, referring to the time during which individuals are infected but not yet infectious. The multi-type version of the birth-death skyline model [12] allows us to distinguish between two types of infected individuals: (i) those who are not yet infectious (typically assigned to a compartment *E*), and (ii) those who are infectious (compartment *I*). Previous work has indicated that phylogenetic tools can estimate the overall infected period (including the exposed and infectious phases), but that it is difficult to estimate the exposed and infectious periods separately [14]. Hence, we run three versions of this analysis, with the infectious period fixed to either 6 months (δ=2), 3 months (δ=4) or 2 months (δ=6) and report the results for each of those setups.

### Data sets

Sampling procedures, strain isolation, sequencing, accession numbers for the sequences, and sequence analysis are described in detail in [3]. In brief, we used the Illumina platform to sequence 68 patients associated with the Bernese outbreak and 46 from the WTK outbreak spanning more than 10 and 36 years, respectively. An alignment of 133 variable positions among 68 isolates from the Bern outbreak [3] and an alignment of 150 variable positions among 30 Californian cases from the WTK outbreak were used [4]. Possible drug resistance conferring mutations as described in [4] were excluded from the alignment. Only one isolate per patient was included and the isolation dates of the strains were used as sampling times.

## Results

### TB in Bern

The Lineage 4 Bernese data set shows positive correlation between genetic divergence and sampling time, but there is little temporal signal (TempEst R^2^=0.05).

The epidemiological parameter estimates we obtain for the Bernese outbreak largely agree among the different model specifications. We estimate that the temporal origin of the Bernese data set was around 1986 with the 95% highest posterior density intervals (HPD) ranging from 1985 – 1988.

Assuming a model without an exposed phase (BDSKY) we estimate an initial high effective reproduction number R_e_ of 4.9 (median, 95% HPD: 2.6-8.1). Around 1991, it declined drastically, staying below the epidemic threshold 1 for the rest of the sampling period (Figure 1). The recovery rate δ is estimated to be 0.2 (median), suggesting an infected period of 5 years. The sampling proportion does not deviate from its prior distribution and is hence estimated at 90%. We estimate that the data set contains one sampled ancestor (95% HPD, 0 – 4), with infected individuals being removed upon sampling with 98% probability (95% HPD, 90-100%). The mean substitution rate for the variant sites is estimated to be 3.9 × 10^−3^ (95% HPD, 2.4 × 10^−3^ − 6.1 × 10^−3^). Table 2 summarizes the median posterior estimates and their 95% highest posterior density (HPD) intervals. Figure 2 shows the maximum clade credibility that was generated from the posterior distribution of trees using TreeAnnotator, which is part of BEAST version 2.4 [10].

**Figure 1:**
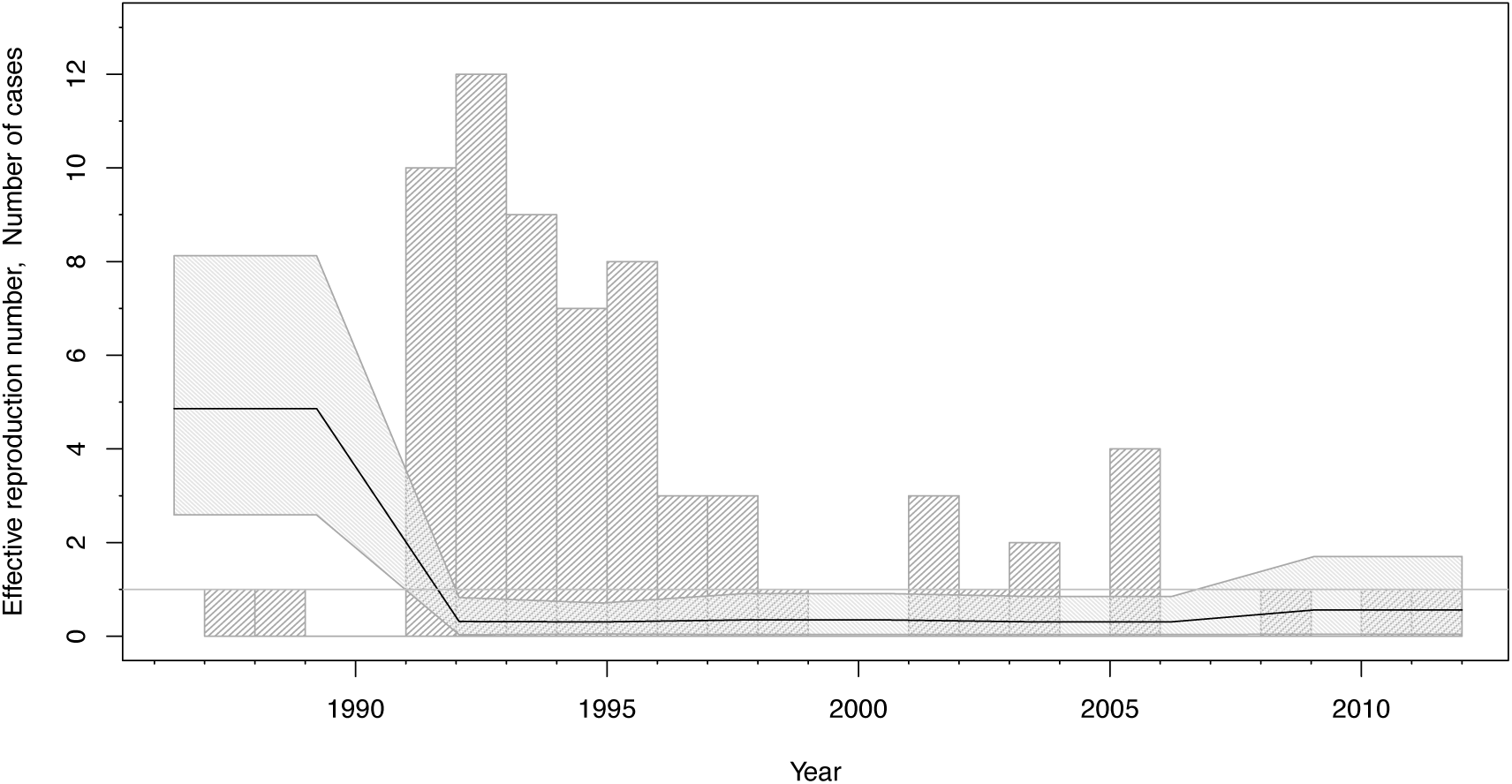
Bern reproduction number through time (bdsky) The median effective reproduction number (black line) with its 951%) highest posterior density (HPD) interval (shaded area). The grey bars display a histogram of the number of cases diagnosed per year.

**Figure 2:**
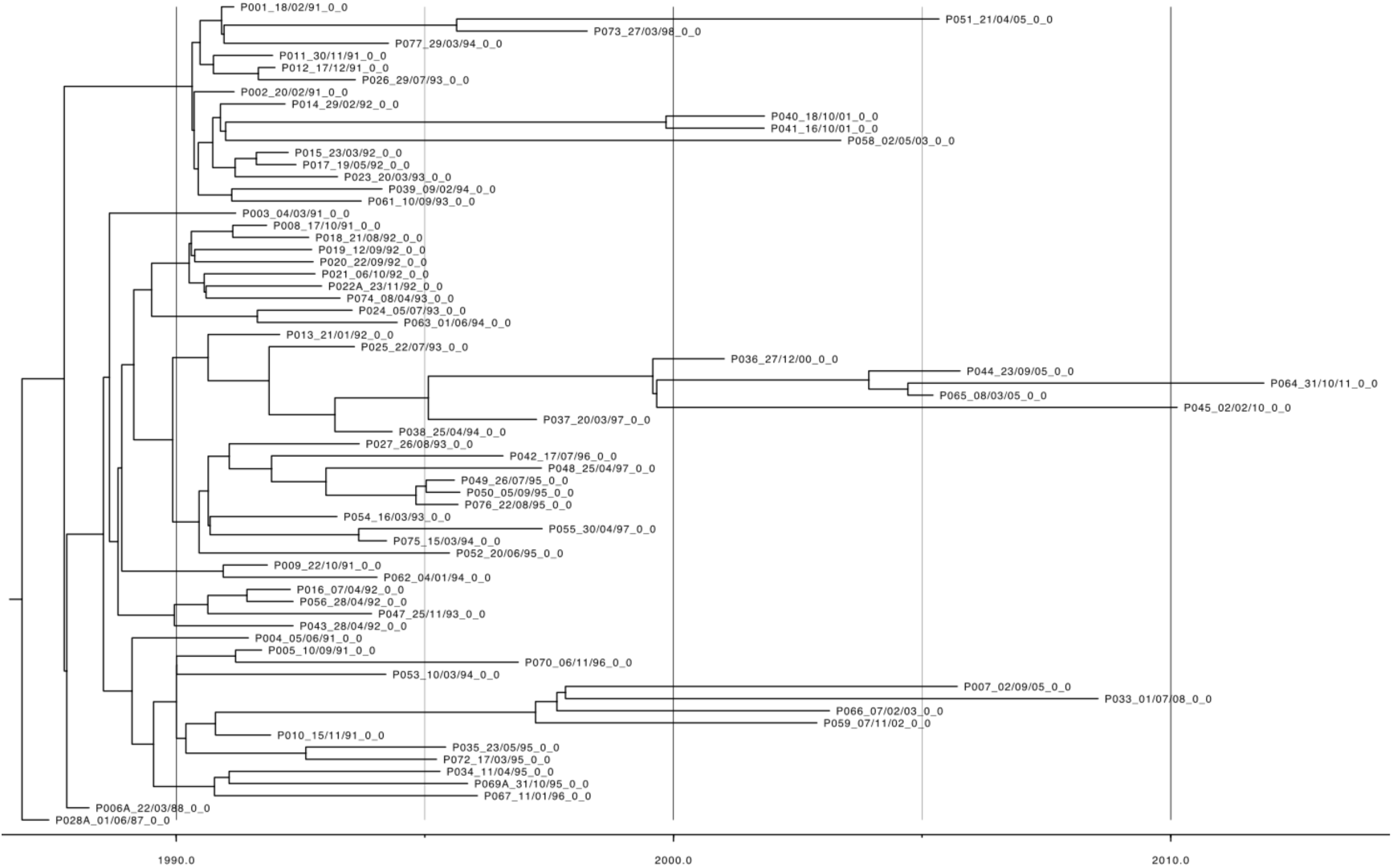
Bern maximum clade credibility tree.

**Table 2.**
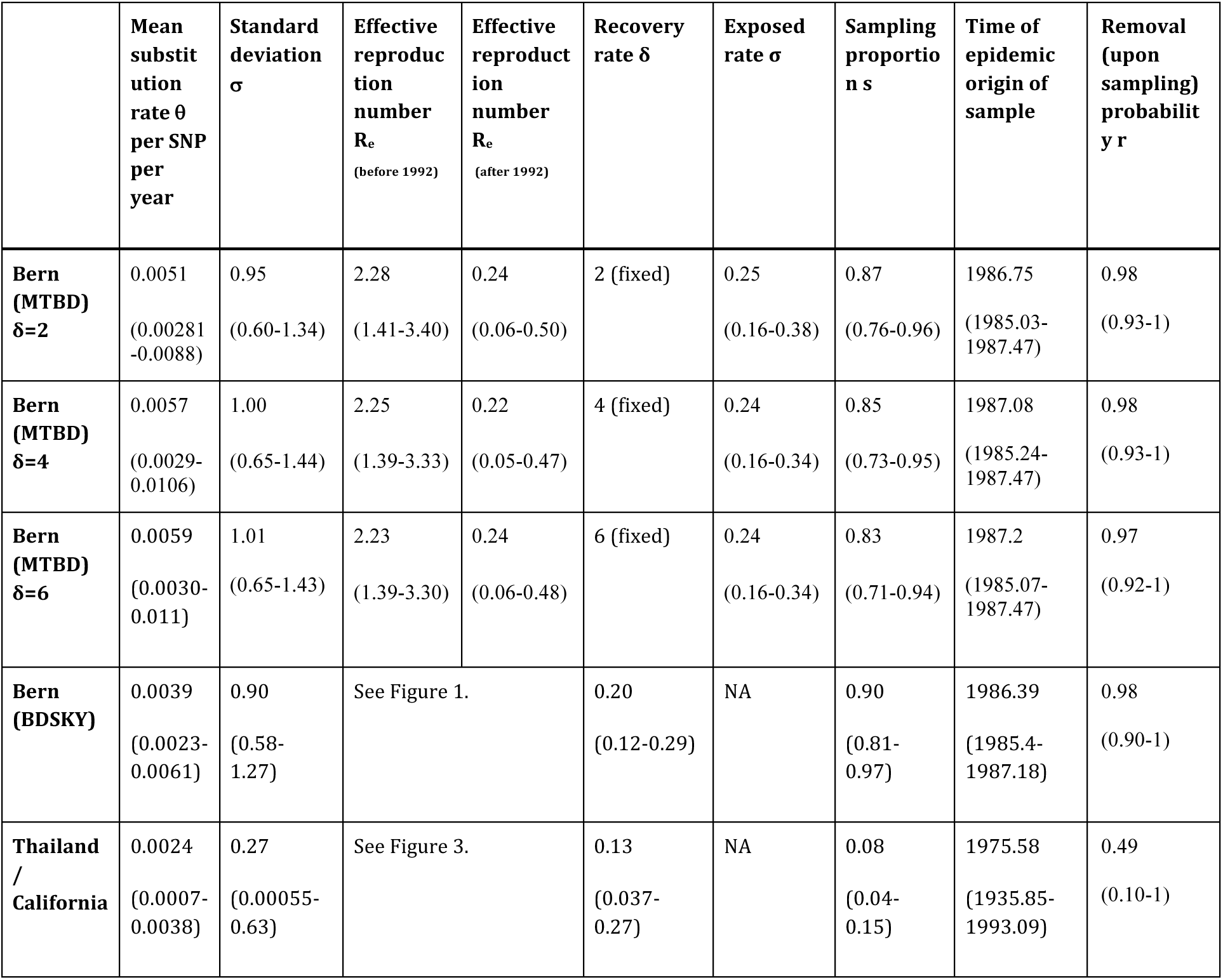

Explicit incorporation of the exposed period in the MTBD model allows us to distinguish the average duration that infected individuals remain infectious. Due to the computational complexity of the model we only allowed one change in the effective reproduction number R_e_ to have occurred in 1992. Before 1992, we estimate median R_e_ values around 2.25 and afterwards the median estimates are significantly below the epidemic threshold 1. Under the MTBD model we fixed the rate δ at which infected individuals become non-infectious to 2, 4 or 6, suggesting an infected period of 6, 3 or 2 months. The median rate σ at which infected individuals become infectious is around 0.25, that is, on average infected individuals became infectious after 4 years in each of the three scenarios. Again, we estimate that the data set contains one sampled ancestor, with infected individuals being removed upon sampling with 98% probability. The mean substitution rate for the variant sites is estimated to be 5.1 × 10^−3^ (95% HPD, 2.8 × 10^−3^ − 8.8 × 10^−3^) when δ=2, 5.7 × 10^−3^ (95% HPD, 2.9 × 10^−3^ − 1.1 × 10^−2^) when δ=4 and 5.9 × 10^−3^ (95% HPD, 3.0 × 10^−3^ − 1.1 × 10^−2^) when δ=6 (Table 2).

### TB in Hmong migrants from Thailand

The Lineage 2 WTK data set shows positive correlation between genetic divergence and sampling time, and a moderate level of temporal signal (TempEst R^2^=0.35).

We analysed this data set under the BDSKY model, and allowed m = 4 intervals to estimate changes in the effective reproduction number R_e_. The temporal origin of this data set is estimated around 1976 with the 95% HPD interval ranging from 1935 – 1993. There is much uncertainty in the estimate of the effective reproduction number R_e_, its median and 95% HPD interval are shown in Figure 3. The rate δ at which infected individuals become non-infectious is estimated to be 0.13 (median), suggesting an infected period of 8 years. The median sampling proportion estimate is 8% (95% HPD, 4 – 15%). We estimate that the data set contains no sampled ancestors (95% HPD, 0 – 2), with the probability to be removed upon sampling r = 64% (95% HPD, 10 – 100%). The mean substitution rate for the variant sites is estimated to be 2.4 × 10^−3^ (95% HPD, 7.2 × 10^−4^ − 3.8 × 10^−3^). Figure 4 shows the maximum clade credibility that was generated from the posterior distribution of trees using TreeAnnotator, which is part of BEAST version 2.4 [10]. Due to the large uncertainty in the BDSKY estimates we did not attempt analysis of the WTK data set under the more complex MTBD model.

**Figure 3.**
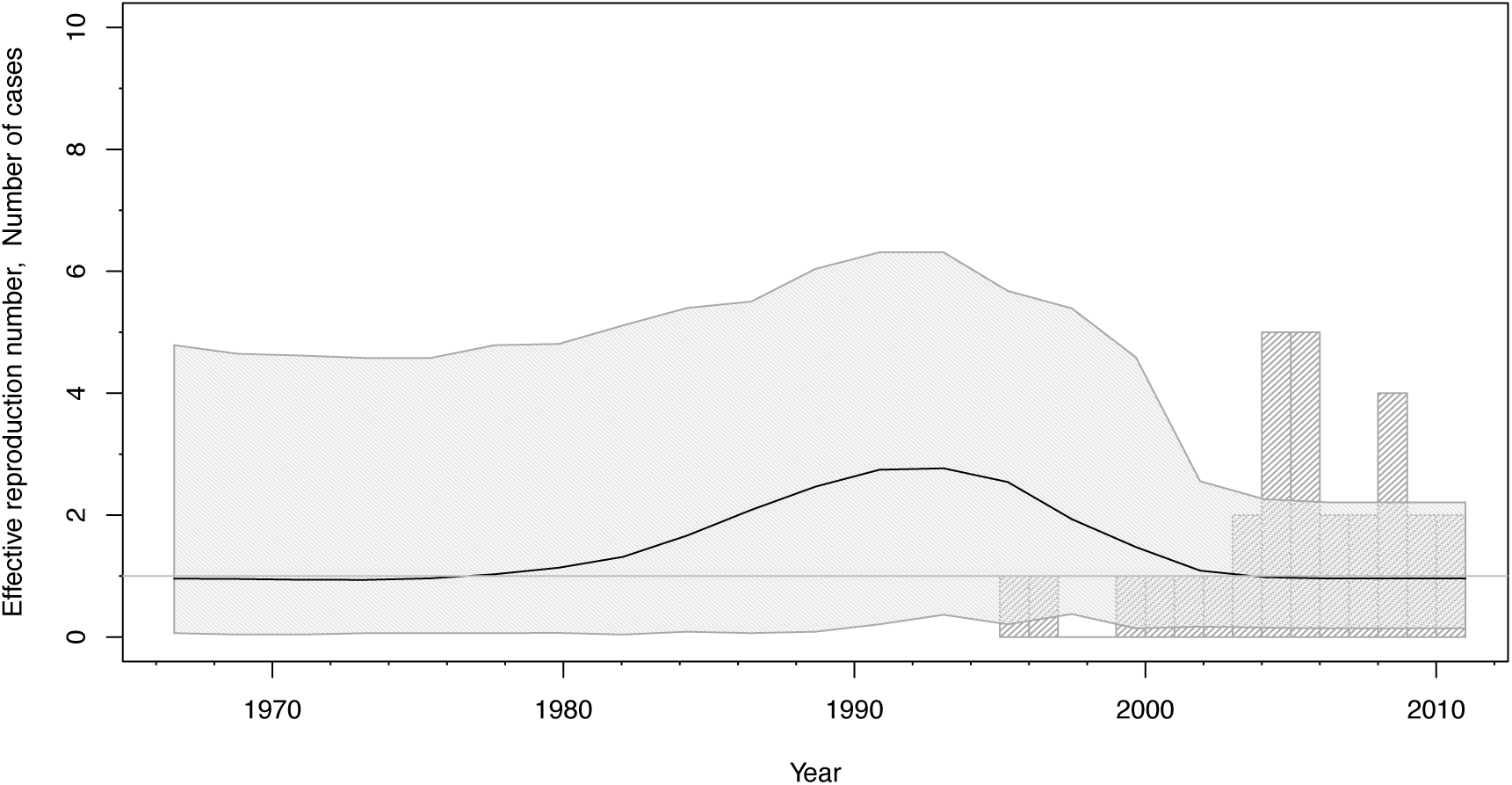
The median effective reproduction number (black line) with its 95% highest posterior density (HPD) interval (shaded area). The grey bars display a histogram of the number of cases diagnosed pier year.

**Figure 4:**
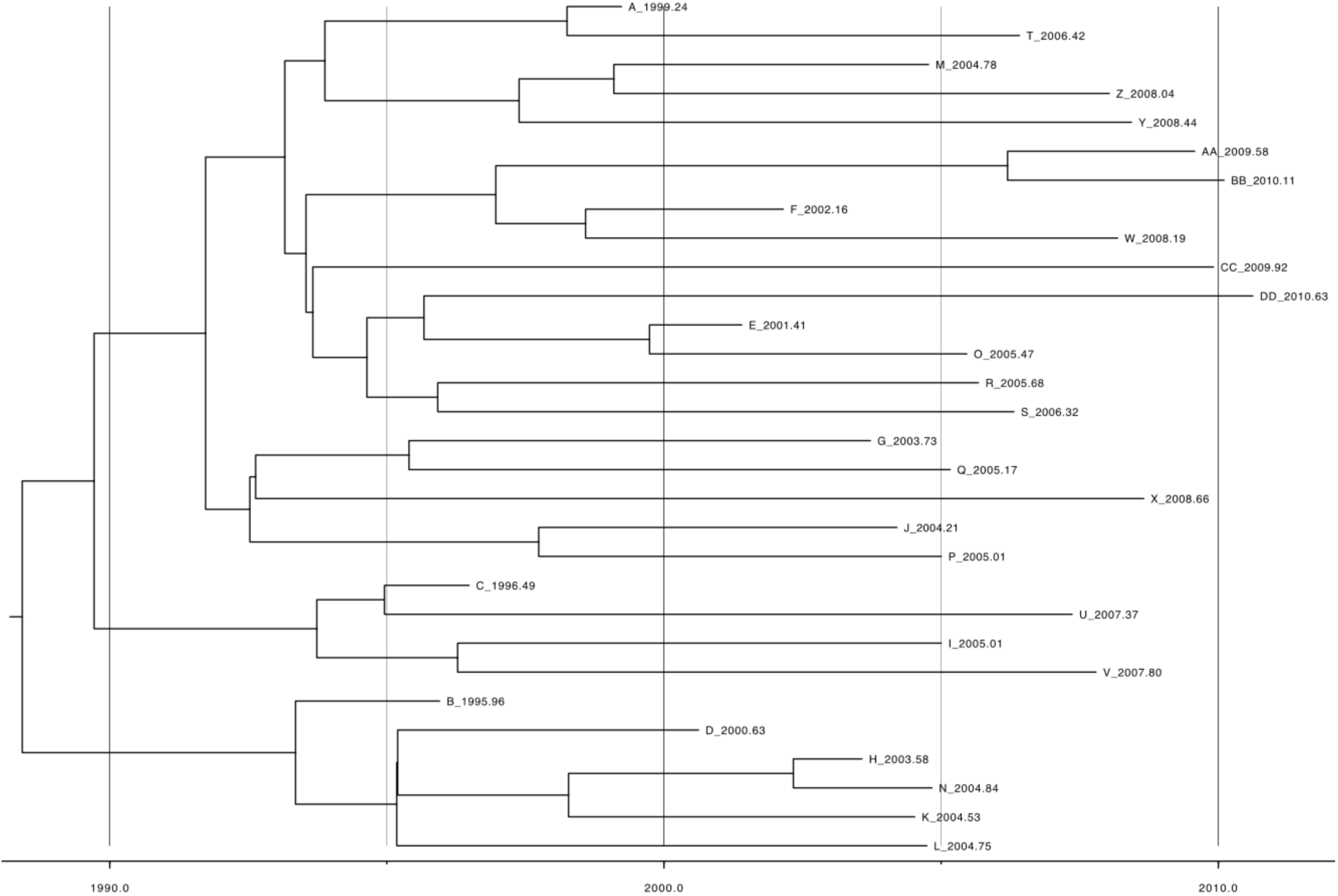
Hmong maximum clade credibility tree.

## Discussion

In this study, we used Bayesian phylodynamic methods to reconstruct the epidemiological dynamics of two *M. tuberculosis* WGS data sets corresponding to two unrelated outbreaks. We quantify the time of the start of the outbreak, and the effective reproductive number through time.

There is much more uncertainty in the epidemiological parameters estimated from the WTK outbreak than the Bern outbreak. The Bernese outbreak is characterized by (i) the outbreak being restricted to the medium size Swiss city Bern and (ii) a large sampling proportion. Indeed, many cases were sampled shortly after infection and subsequent cases have been recovered by a SNP screening assay and targeted WGS, such that an estimated 90% of secondary (i.e. infectious) cases linked to this outbreak are included in the data set. This high sampling proportion and the geographical containment of the Bernese outbreak are the likely cause of the higher confidence of the estimates obtained for this dataset, because the sampling times of sequences in a densely-sampled outbreak are very informative for the age of an outbreak, and thus allow to time-calibrate the phylogeny and quantify transmission and recovery rates. On the other hand, the WTK data set has roughly half the number of samples and a much lower sampling proportion of 9% (median estimate). Furthermore, although the outbreak started in Thailand, our WTK data set consists entirely of cases imported into California. This sparse outbreak sample does not contain much information regarding the age of the outbreak, resulting in much uncertainty in the epidemiological estimates.

Our analyses were conducted using phylodynamic methods implemented in the Bayesian MCMC framework BEAST version 2.4 [15], which means that we are estimating so-called time-trees using molecular clock models. Before using such models one should explore the temporal signal in sequence alignments, which can be done using TempEst [9]. While both data sets exhibit a positive correlation between genetic divergence and sampling time, there is a moderate level of temporal signal only in the WTK data set (R^2^=0.35). After scaling to account for SNP alignment, we obtain a median evolutionary rate of 6.7 × 10^−8^ for the WTK outbreak. The WTK data set belongs to Lineage 2, for which Duchene et al. [16] were unable to reliably determine the evolutionary rate. As outbreak data sets are often not suitable for mutation rate estimation this estimate should be taken with a grain of salt. For a robust estimate one would want to collect longitudinal data over a longer time period [16].

There is little temporal signal in the Lineage 4 Bernese data set (R^2^=0.05), which explains the uncertainty in our clock rate estimates of the Bernese outbreak. Our results show that the estimated time of the epidemic origin and the epidemiological parameters are robust to the differing clock rate estimates, see Table 2.

We hypothesize that the two data sets are an example of the time dependency of molecular rate estimates [17]: the estimates of the evolutionary rate for the Bernese outbreak represent a high short-term rate of evolution, whereas due to the delayed sampling, the WTK estimate is a low longer-term mutation rate of evolution. Hence, our evolutionary rate estimates are not suitable for comparison between the two *M. tuberculosis* lineages.

Our phylodynamic analyses allowed us to estimate the temporal dynamics of the Bernese outbreak. Despite the fact that the sampling dates range from 1987 to 2011, our results support the hypothesis that the epidemic peaked around 1990 [3]. This indicates that the peak of the outbreak occurred several years before it was detected. Indeed, most of the transmission events likely occurred between 1990 and 1991, although the majority of cases was only reported in 1993 [3]. This refutes the previous hypothesis that disease would have occurred shortly after infection, with short latent periods [18], due to the population characteristics in the affected population of the Bernese outbreak (homeless, substance abusers, etc.).

Both models employed for analysis of the Bernese outbreak (BDSKY and MTBD) suggest that the average infected period lasted about 4-5 years. While in BDSKY the infected period is equivalent to the infectious period, the infected period in the MTBD model is the sum of the infectious and exposed periods. In the latter we assume an infectious period of 2, 3 or 6 months [19], and in each of those cases the exposed period is robustly estimated around 4 years. While this means that both models agree on the overall infected period to last around 4-5 years, we know that MTBD is the more realistic model. Hence, we conclude that an – on average – infected patient in the Bern outbreak was in the latent/exposed stage of the disease for about 4 years before becoming infectious and consequently being diagnosed and treated shortly after [19].

For the WTK outbreak, we estimated an infectious period of eight years, which is significantly higher than the infectious period estimated for the Bernese outbreak (p-value < 2.2 × 10^−16^). This may be due to a delay in sampling and treatment, due to the sampling having taken place in California only, such that patients were likely sick and infectious for longer. Furthermore, while the WTK outbreak was caused by an MDR strain, the Bernese outbreak was caused by a sensitive strain. Resistance is a likely cause of delayed treatment success [20].

Our study shows that phylodynamic analysis of WGS data can shed light on the temporal dynamics of tuberculosis outbreaks. Analysis of the Bernese outbreak has revealed that even when there is little temporal signal, we can robustly estimate epidemiological parameters if the sampling proportion is large. Conversely, in the WTK outbreak there is much uncertainty in the epidemiological parameter estimates despite a moderate temporal signal. This may be due to a difference in transmission dynamics in Thailand versus California as well as the fact that the epidemic peak likely occurred before the first samples were taken.

Overall, we believe that real time outbreak WGS together with phylodynamic methods will improve future outbreak investigation as phylodynamic analysis can shed light on the timing of the epidemic origin and transmission dynamics through time.

## Acknowledgements

DK gratefully acknowledges support from the ETH Zürich Postdoctoral Fellowship Program and the Marie Curie Actions for People COFUND, and the Swiss National Science Foundation (SNSF) for generously funding her research with a Marie Heim-Vögtlin fellowship. TS is supported in part by the European Research Council under the Seventh Framework Programme of the European Commission (PhyPD grant 335529). This work was further supported by the Swiss National Science Foundation (grant 310030_166687 to S.G.), the European Research Council (309540-EVODRTB to S.G.), and SystemsX.ch. The work on the tuberculosis outbreak investigations was supported by a grant from the Bernese Lung Association (LF). The funders had no role in study design, data collection and analysis, decision to publish, or preparation of the manuscript.

